# Autosomal dominant multiple pterygium syndrome is caused by mutations in *MYH3*

**DOI:** 10.1101/017434

**Authors:** Jessica X. Chong, Lindsay C. Burrage, Anita E. Beck, Colby T. Marvin, Margaret J. McMillin, Kathryn M. Shively, Tanya M. Harrell, Kati J. Buckingham, Carlos A. Bacino, Mahim Jain, Yasemin Alanay, Susan A. Berry, John C. Carey, Richard A. Gibbs, Brendan H. Lee, Deborah Krakow, Jay Shendure, Deborah A. Nickerson, University of Washington Center for Mendelian Genomics, Michael J. Bamshad

**Affiliations:** Division of Genetic Medicine, Department of Pediatrics, University of Washington, Seattle, WA 98195, USA; Department of Molecular and Human Genetics, Baylor College of Medicine, One Baylor Plaza, Suite R814, Houston, TX, 77030-3411, USA; Division of Genetic Medicine, Seattle Children’s Hospital, Seattle, WA 98105, USA; Pediatric Genetics, Department of Pediatrics, Acibadem University School of Medicine, istanbul, Turkey 34752; Division of Genetics and Metabolism, Department of Pediatrics, University of Minnesota, Minneapolis, MN, 55455, USA; Department of Pediatrics, University of Utah, Salt Lake City, UT 84108, USA; Department of Molecular and Human Genetics, Baylor College of Medicine, Houston, TX 77030, USA; Human Genome Sequencing Center, Baylor College of Medicine, Houston, TX 77030, USA; Department of Human Genetics and Department of Orthopedic Surgery, University of California Los Angeles, California, USA 90048; Department of Genome Sciences, University of Washington, Seattle, WA 98195, USA

**Keywords:** exome sequencing, Mendelian disease, congenital contractures, distal arthrogryposis, multiple pterygium syndrome

## Abstract

Multiple pterygium syndromes (MPS) are a phenotypically and genetically heterogeneous group of rare Mendelian conditions characterized by multiple pterygia, scoliosis and congenital contractures of the limbs. MPS typically segregates as an autosomal recessive disorder but rare instances of autosomal dominant transmission have been reported. While several mutations causing recessive MPS have been identified, the genetic basis of dominant MPS remains unknown. We identified four families with dominantly transmitted MPS characterized by pterygia, camptodactyly of the hands, vertebral fusions, and scoliosis. Exome sequencing identified predicted protein-altering mutations in embryonic myosin heavy chain (*MYH3*) in three families. *MYH3* mutations underlie distal arthrogryposis types 1, 2A and 2B, but all mutations reported to date occur in the head and neck domains. In contrast, two of the mutations found to cause MPS occurred in the tail domain. The phenotypic overlap among persons with MPS coupled with physical findings distinct from other conditions caused by mutations in *MYH3*, suggests that the developmental mechanism underlying MPS differs from other conditions and / or that certain functions of embryonic myosin may be perturbed by disruption of specific residues / domains. Moreover, the vertebral fusions in persons with MPS coupled with evidence of *MYH3* expression in bone suggests that embryonic myosin plays a previously unknown role in skeletal development.

---

Multiple pterygium syndromes (MPS) are a phenotypically and genetically heterogeneous gxzroup of rare Mendelian conditions characterized by multiple pterygia, scoliosis and congenital contractures of the limbs. Most often, MPS occurs as a simplex case, and of reported multiplex families, the majority consist of multiple affected siblings born to unaffected parents, consistent with inheritance in an autosomal recessive pattern^1^. In very rare instances, MPS has been transmitted from an affected parent to an affected child, indicative of autosomal dominant transmission^2-5^. In 1996, we revised the classification of distal arthrogryposis (DA) syndromes and categorized several additional conditions^6^, including autosomal dominant MPS, as DA syndromes because their clinical features overlapped with DA type 1 (DA1; MIM 108120) and Freeman-Sheldon syndrome or DA2A (MIM 193700). Autosomal dominant MPS, or DA type 8 (DA8; MIM 178110), was one of the conditions added to the DA classification based on the phenotypic features of four reported families in which pterygia, camptodactyly of the hands, vertebral fusions, and scoliosis, were transmitted from parent-to-child^2-5^.

Over the past decade, three as of yet unreported families with multiple persons who had clinical characteristics consistent with the diagnosis of DA8 and evidence of parent-to-child transmission were referred to our research program on DA syndromes (Table 1; Figures 1 and 2; Figure S1). To identify the gene(s) harboring mutations underlying DA8, we initially screened by Sanger sequencing the proband of each family for mutations in genes known to underlie lethal MPS (MIM 253290) and non-lethal Escobar-variant autosomal recessive MPS (MIM 265000) including *CHRNG*^7,8^ (MIM 100730), *CHRND*^9^ (MIM 100720), *CHRNA1*^9^ (MIM 100690) as well as *IRF6*^10^ (MIM 607199), mutations in which cause popliteal pterygium syndrome (MIM 119500). No pathogenic mutations were found in any of these candidate genes. Next, we preformed exome sequencing on seven individuals in family A, three individuals in family B, and two affected individuals (III-5 and V-2) in family C (Table 1; Figures 1 and 2; Figure S1).All studies were approved by the institutional review boards of the University of Washington and Seattle Children’s Hospital; and informed consent was obtained from each participant.

**Figure 1 (not included).**
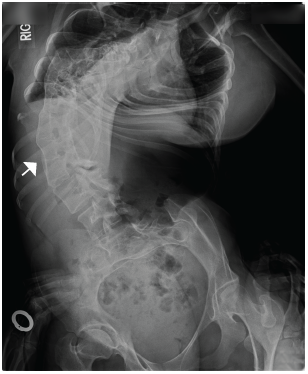
Phenotypic Characteristics of Individuals with Distal Arthrogryposis type 8. Five individuals affected with DA8; all individuals shown have *MYH3* mutations. Note the downslanting palpebral fissures, ptosis, camptodactyly of the fingers, scoliosis, short stature, and neck webbing. Case identifiers for the individuals shown in this figure correspond to those in Table 1, where there is a detailed description of the phenotype of each affected individual. Figure S1 provides a pedigree of each family with DA8.

**Table 1.**
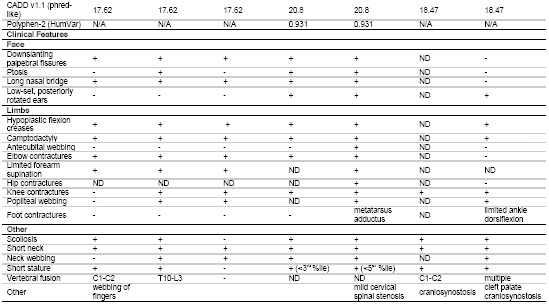
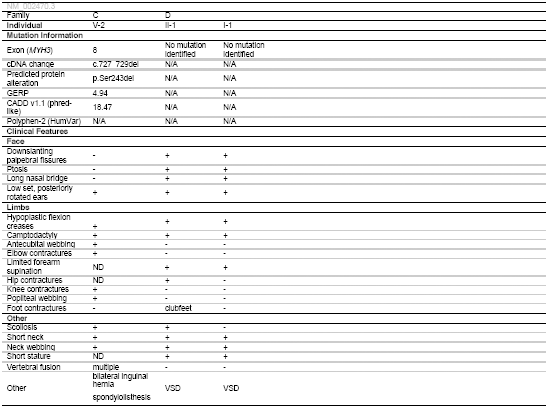
Mutations and Clinical Findings of Individuals with Distal Arthrogryposis type 8. This table provides a summary of clinical features of affected individuals from families in which mutations in *MYH3* causing DA8 were identified and one family in which no mutation was identified. Clinical characteristics listed in table are primarily features that delineate DA8. Plus (+) indicates presence of a finding, minus (-) indicates absence of a finding. * = described per report. ND = no data were available. N/A = not applicable. GERP = Genomic Evolutionary Rate Profiling. CADD = Combined Annotation Dependent Depletion, VSD = ventricular septal defect. cDNA positions provided as named by the HGVS MutNomen web tool relative to NM_002470.3.

In brief, one microgram of genomic DNA was subjected to a series of shotgun library construction steps, including fragmentation through acoustic sonication (Covaris), end-polishing (NEBNext End Repair Module), A-tailing (NEBNext dA-Tailing Module), and PCR amplification with ligation of 8 bp barcoded sequencing adaptors (Enzymatics Ultrapure T4 Ligase) for multiplexing. One microgram of barcoded shotgun library was hybridized to capture probes targeting ∼36.5 Mb of coding exons (Roche Nimblegen SeqCap EZ Human Exome Library v2.0). Library quality was determined by examining molecular weight distribution and sample concentration (Agilent Bioanalyzer). Pooled, barcoded libraries were sequenced via paired-end 50 bp reads with an 8 bp barcode read on Illumina HiSeq sequencers.

Demultiplexed BAM files were aligned to a human reference (hg19) using the Burrows-Wheeler Aligner (BWA) 0.6.2. Read data from a flow-cell lane were treated independently for alignment and QC purposes in instances where the merging of data from multiple lanes was required. All aligned read data were subjected to: (1) removal of duplicate reads (Picard MarkDuplicates v1.70) (2) indel realignment (GATK IndelRealigner v1.6-11-g3b2fab9); and (3) base quality recalibration (GATK TableRecalibration v1.6-11-g3b2fab9). Single nucleotide variant (SNV) detection and genotyping were performed using GATK UnifiedGenotyper (v1.6-11-g3b2fab9). SNV data for each sample were formatted (variant call format [VCF]) as “raw” calls that contained individual genotype data for one or multiple samples, and flagged using the filtration walker (GATK) to mark sites that were of lower quality and potential false positives (e.g. strand bias > -0.1, quality scores (Q50), allelic imbalance (ABHet≥0.75), long homopolymer runs (HRun>3), and/or low quality by depth (QD<5).

Because DA8 is extremely rare, with only four families reported to date^2-5^, we excluded SNVs with an alternative allele frequency >0.0001 in any population in the NHLBI Exome Sequencing Project Exome Variant Server (ESP6500/EVS), 1000 Genomes, or Exome Aggregation Consortium (ExAC v1.0) browser, or in an internal exome database of ∼700 exomes. Additionally, SNVs that were flagged as low quality or potential false positives (quality score ≤ 30, long homopolymer run > 5, low quality by depth < 5, within a cluster of SNPs) were also excluded from analysis. Copy number variant (CNV) calls were generated from exome data with CoNIFER^11^. Variants were annotated with the SeattleSeq138 Annotation Server, and SNVs for which the only functional prediction label was intergenic, coding-synonymous, utr, near-gene, or intron were excluded. Individual genotypes with depth<6 or genotype quality<20 were treated as missing in the analysis.

To exclude known causes of multiple pterygium syndromes and well-known conditions that include pterygia, we first assessed the exome data for pathogenic variants in the genes previously screened by Sanger sequencing (*IRF6, CHRNG, CHRNA1*, and *CHRND*) as well as in *RAPSN*^9,12^ (MIM 601592) and *DOK7*^13^ (MIM 610285). No SNVs or CNVs in these genes segregated with DA8 in any of the three families tested. Next, we looked for novel and/or rare SNVs and CNVs in the same gene shared among affected probands. No genes with rare or novel variants were shared among all three families. Affected individuals in Families A and B were also screened for larger copy number variations (CNVs) by comparative genome-wide array genomic hybridization on the Illumina Infinium HumanCore-24 chip. No shared or overlapping CNVs were identified. Subsequently, we loosened our filtering criteria to include variants in genes that were shared by two out of three families and identified three candidate genes (*SHPRH* [MIM 608048], *SLC5A9, MYH3*) in families A and B.

The known functions of *SHPRH* and *SLC5A9* appeared to be unrelated to the musculoskeletal phenotypes observed in all three families: *SHPRH* is involved in DNA repair and maintenance of genomic stability while *SLC5A9* encodes a sugar transporter primarily expressed in small intestine and kidney. In contrast, mutations in *MYH3* frequently cause other forms of distal arthrogryposis, specifically DA2A and DA2B^14^ (MIM 601680), and more rarely, DA1^15,16^. In addition to congenital contractures, persons with DA2A and DA2B variably have short stature, scoliosis, and infrequently pterygia of the neck^17^. Accordingly, we considered *MYH3* to be the most compelling candidate gene in these families.

In *MYH3* (NM_002470.3), we specifically discovered a c.3214_3216dup, [p.(Asn1072dup)] variant in family A and a c.3224A>C [p.(Gln1075Pro)] variant in family B. Both variants were subsequently validated via Sanger sequencing and found to segregate with only the affected persons in each family (Figure 2). Both variants affect highly conserved amino acid residues, with identical Genomic Evolutionary Rate Profiling (GERP) scores of 5.64, and are predicted to be deleterious by multiple methods including a Combined Annotation Dependent Depletion (CADD^18^) score of 17.62 for p.(Asn1072dup) and 20.8 for p.(Gln1075Pro). Moreover, neither variant was found in >71,000 control exomes comprised of the ESP6500, 1000 Genomes phase 1 (Nov 2010 release), internal databases (>1,400 chromosomes), and ExAC (October 20, 2014 release).

**Figure 2.**
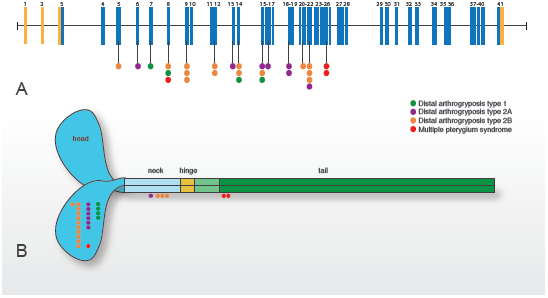
Radiograph of severe scoliosis and vertebral fusion observed in a person (AIII-2, Table 1) with Distal Arthrogryposis type 8. Severe, convex rightward, rotatory scoliosis of the thoracic and lumbar spine. Fusion of vertebral bodies from T10 to L3 is indicated by the arrow.

Independent of the effort at the University of Washington to identify a gene associated with DA8, investigators at Baylor College of Medicine had identified a 5-generation family in which 10 persons were reported to be variably affected with a dominantly-inherited condition characterized by short stature, scoliosis, multiple vertebral fusions, camptodactyly of the fingers, and multiple pterygia (Table 1; Figure S1), consistent with the diagnosis of DA8. Four individuals from this family were enrolled in a research protocol approved by the institutional review board at Baylor College of Medicine. Informed consent was obtained from each of these individuals prior to enrollment and prior to initiation of research studies. Whole exome sequencing was performed with DNA obtained from peripheral blood monocytes from the proband (Individual IV-3, Figure S1) and her father (Individual III-5, Figure S1) using the VCRome 2.1 target capture reagents (Roche Nimblegen)^19^. Sequencing was conducted on Illumina HiSeq2000 as previously described^19^. Alignment, variant calling and annotation was completed using the *Mercury* pipeline^20^. Comparison of exome sequence data from the proband and her affected father (Table 1; Figure S1) identified 46 shared rare variants (i.e., frequency<0.0001 in ESP6500, 1000 Genomes phase 1 (Nov 2010 release), and ExAC (October 20, 2014 release)). No variants were identified in *SHPRH* or *SLC5A9*, however, one novel variant,c.727_729del, (p.Ser243del) affected a highly conserved amino acid residue of *MYH3* (GERP score of 4.94) and was predicted to be highly deleterious (CADD v1.1 score of 18.47). This variant was validated by Sanger sequencing and segregated in all of the affected individuals for whom DNA was available for testing. This variant has not been observed in ESP6500, ExAC (Version 0.3, January 13, 2015 release) and 1000 Genomes (October 2014 release).

In 2006, we reported that mutations in *MYH3* cause DA2A and DA2B^14^. Nevertheless, *MYH3* was never considered a high priority candidate gene in individuals with DA8 because affected individuals had multiple pterygia of the limbs, severe scoliosis, and vertebral fusions and did not have contractures of the facial muscles. More recently, we and others, identified mutations in *MYH3* in persons with DA1 who had no facial contractures^15,16^ and persons with DA2B who had vertebral fusions and severe congenital scoliosis^21^ indicating that the phenotypic spectrum associated with *MYH3* variants is broader than originally considered. This inference is now supported by our observation that variants in *MYH3* cause DA8 and a wide range of phenotypic abnormalities of connective tissue including congenital contractures, multiple pterygia, and bony fusions.

*MYH3* encodes embryonic myosin heavy chain, a skeletal muscle myosin composed of a globular motor domain (amino acid residues ∼1-779) attached by a short neck (∼779-840) and hinge (∼840) region to a long coiled-coil rod domain (∼840-1940). The majority of the rod region comprises the myosin tail domain (amino acid residues ∼1070-1940). Embryonic myosin exists as a dimer in which the tail domains are intertwined (Figure 3). Hundreds of myosin dimers assemble with one another and other proteins to form the thick filaments of the sarcomere of myofibrils, the latter of which represents the subcellular contractile apparatus of cardiac and skeletal muscle cells.

**Figure 3.**
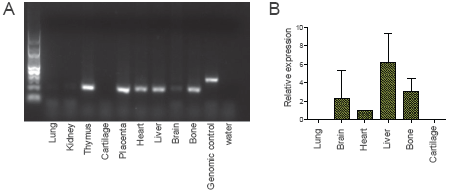
Genomic Model of *MYH3* and stylized structure of Embryonic Myosin Illustrating spectrum of mutations that cause Distal Arthrogryposis Syndromes. MYH3 is composed of 41 exons, 39 of which are protein-coding (blue) and two of which are non-coding (orange). A) Lines with attached dots indicate the approximate locations of mutations in *MYH3* that cause DA1 (green), DA2A (purple), DA2B (orange), and DA8 (red). The color of each dot reflects diagnosis. B) Schematic of an embryonic myosin dimer. The approximate locations of mutations relative to the myosin domains are indicated by colored circles. All mutations causing DA1, DA2A, and DA2B are located in the motor and neck of MYH3, while the two mutations causing DA8 are the only mutations located in the tail. (Both variants in the tail of MYH3 [c.4934A>C; p.(Asp1622Ala) and c.2979C>T; p.(Ala1637Val)] reported by Toydemir et al. 2006^14^ are now thought to be polymorphisms based on additional family information and frequencies in large control population databases.)

All of the mutations known to cause DA1, DA2A, and DA2B^16^ affect amino acid residues in the head and neck domains of embryonic myosin whereas two of the three mutations that cause DA8 occur in the tail domain. The clinical characteristics of the individuals with DA8 and a mutation in the tail domain were more similar to one another compared to the features of persons with a mutation in the head domain. This observation is limited, of course, by the small number of both families and individuals with DA8 caused by *MYH3* mutations in our dataset. Nevertheless, speculation on domain-specific or at least domain-predominant phenotypes has been reported previously for mutations in both muscle and non-muscle myosins. For example, mutations in the head domain of *MYH9* (MIM 160775) cause severe thrombocytopenia and are associated with nephritis and deafness, while mutations in the tail domain cause less severe thrombocytopenia, with median platelet counts more than two times as high as those associated with head domain mutations^22^. Similarly, mutations throughout *MYH7* (MIM 160760) can cause hypertrophic (MIM 192600) and/or dilated cardiomyopathy (MIM 613426) while skeletal myopathies (e.g., myosin storage myopathy [MIM 608358] and Laing distal myopathy [MIM 160500]) are caused almost exclusively by mutations in the tail domain of *MYH7*^23^. Moreover, mutations in the tail of *MYH7* disrupt stability or self-assembly of sarcomere filaments or protein-protein interactions mediated through the tail, whereas mutations in the motor appear to alter the ATPase and actin-binding properties of myosin^24^. This precedent of domain-specific phenotypic differences associated with mutations in other myosin genes suggest that similar genotype-phenotype relationships might exist in *MYH3*.

Skeletal abnormalities have not commonly been observed, or at least reported, in persons with DA1, DA2A, or DA2B and mutations in *MYH3*. Yet, skeletal defects were found in each of the three DA8 families with an *MYH3* mutation that we studied. The most common defects observed were vertebral abnormalities including hemivertebrae and vertebral fusions of C1 and C2 or vertebrae of the thoracolumbar spine (Figure 2). A similar range of vertebral abnormalities has been reported previously^2-5^ in families with dominantly inherited MPS—as well as both carpal and tarsal fusions. In addition, two persons in Family C had craniosynostosis. These observations suggest that embryonic myosin plays a role in skeletal development that is perturbed by the mutations in *MYH3* that cause DA8. Such a role for embryonic myosin would not be unprecedented as other proteins of the contractile apparatus of skeletal muscle, such as troponin I type 2 (skeletal, fast) (*TNNI2*, MIM 191043), are expressed in osteoblasts and chondrocytes in long bone growth plates and play a role in skeletal development^25^. Moreover, multiple myosins have been reported to interact with *Runx2* in rat osteoblasts and osteoblast differentiation *in vitro* was associated with the expression of myosin^26^.

To investigate this possibility further, we used primers specific for *MYH3* cDNA that we had previously developed^27^ to test for *MYH3* expression in a variety of human fetal tissues. As expected, *MYH3* was expressed in thymus, placenta, heart, and liver (Figure 4A). However, whereas *MYH3* expression was not detectable in cartilage, it was expressed in bone (Figure 4A). Next, to determine the relative expression levels of *MYH3,* we conducted a quantitative PCR assay and confirmed *MYH3* expression in bone at levels comparable to several other tissues, including brain, with known *MYH3* expression^28^ (Figure 4B). These preliminary findings suggest that the bony fusions in individuals with DA8 disrupt a previously unknown role of embryonic myosin in vertebral development, and perhaps in skeletal development in general.

Five simplex cases of MPS referred to our center had clinical characteristics that overlapped with DA8 and in several cases were indistinguishable from DA8. Exome sequencing did not reveal *MYH3* mutations in any of these cases, but three of them were explained by mutations in *CHRNG*. Specifically, two persons were compound heterozygous (NM_005199.4 c.[753_754delCT];[805+1G>A] and c.[459dupA];[1080dupC;1082dupC]) and one was homozygous (c.459dupA) for *CHRNG* mutations. Two of these mutations, c.753_754delCT and c.459dupA, have been reported in multiple individuals with recessively inherited MPS^1,8^ and have been described as “recurrent.” Yet, both of these mutations are in EVS and ExAC at frequencies as high as 0.002 (e.g. c.753_754delCT in European-Americans in EVS), suggesting that standing rather than *de no*vo variation contributes to the relatively high incidence of MPS caused by mutations in *CHRNG*. Moreover, c.459dupA occurs on an identical ∼700 kb haplotype in each of the apparently unrelated persons with MPS whom we tested. Therefore,the c.459dupA mutation likely arose many generations ago, and previous attempts did not successfully identify the shared haplotype because of its small size^1^.

Our findings collectively suggest that features such as scoliosis and pterygia observed in individuals with DA8 likely have a different underlying molecular mechanism(s) than in recessive forms of MPS caused by mutations in the genes, *CHRNG, CHRND, CHRNA1*^9^, encoding subunits of the skeletal muscle nicotinic acetylcholine receptor (AChR). AChR is required for normal muscle development and muscle biopsies from individuals with recessive MPS have demonstrated abnormal distribution of AChR and acetylcholinesterase^29^. Since AChR is necessary for proper organization and establishment of the neuromuscular junction^30^ as well as muscle and synaptic maturation^31^, it has been suggested that many features of recessive MPS such as the joint contractures and pterygia are ultimately caused by fetal akinesia^7^ due to abnormal muscle and synaptic development. In contrast, mutations in *MYH3* that cause DA2A/B are hypothesized to alter contractile function of skeletal myofibers during development, resulting in altered movement in utero and congenital contractures. Consistent with this hypothesis, we have recently demonstrated that the p.(Arg672Cys) variant in *MYH3*, the most common cause of DA2A, causes reduced contractile force and prolonged relaxation of human skeletal myofibers^27^. Together, these findings suggest that pterygia form via different developmental mechanisms in persons with DA8 versus autosomal recessive MPS.

We failed to identify the cause of dominantly inherited MPS in one family originally diagnosed with DA8 (Family D in Table 1 and Figure S1). In retrospect, the phenotype in this family differs from that found in the other DA8 families. Specifically, neither of the affected persons in this family had popliteal or antecubital ptergyia and both persons lacked vertebral fusions. Additionally, each individual had a ventriculoseptal defect. Accordingly, while the lack of an *MYH3* mutation in this family could be interpreted as evidence that DA8 is genetically heterogeneous, the alternative is that the phenotype in this family is distinctive enough to prompt delineation of a separate Mendelian condition.

In summary, we used exome sequencing to discover that mutations in *MYH3* cause DA8. The phenotypic overlap among persons with DA8 coupled with physical findings distinct from other DA conditions caused by mutations in *MYH3*, suggests that the developmental mechanism underlying DA8 differs from other DA conditions or that certain functions of embryonic myosin may be perturbed by disruption of specific residues / domains.

## Supplemental Data

Supplemental Data includes 1 figure.

## Acknowledgements

We thank the families for their participation and support; Christa Poel, Karynne Patterson, Jennifer Chin, Yuqing Chen, Alyssa Tran, Philippe Campeau, and James Lu for technical assistance and Alice Ward Racca for helpful comments. Our work was supported in part by grants from the National Institutes of Health / National Human Genome Research Institute and the National Heart, Lung and Blood Institute (1U54HG006493 to M.B., D.N., and J.S.; 1RC2HG005608 to M.B., D.N., and J.S.; 5R000HG004316 to H.K.T.), National Institute of Child Health and Human Development (1R01HD048895 to M.J.B.), the Life Sciences Discovery Fund (2065508 and 0905001), and the Washington Research Foundation. This work was also supported by the Baylor College of Medicine Intellectual and Developmental Disabilities Research Center (HD024064) from the Eunice Kennedy Shriver National Institute Of Child Health & Human Development, NIH U54HG003273 (R.A.G.), NIH U54 U54HG006542 (R.A.G.), NIH T32 GM07526 (M.J. and L.C.B.), and NICHD P01 HD070394 (B.L.). L.C.B. was also supported by the Genzyme/ACMG Foundation for Genetic and Genomic Medicine Medical Genetics Training Award in Clinical Biochemical Genetics, the National Urea Cycle Disorders Foundation Fellowship, and a fellowship from the Urea Cycle Disorders Consortium (UCDC; U54HD061221), which is a part of the National Institutes of Health (NIH) Rare Disease Clinical Research Network (RDCRN), supported through collaboration between the Office of Rare Diseases Research (ORDR), the National Center for Advancing Translational Science (NCATS and the Eunice Kennedy Shriver National Institute of Child Health and Human Development (NICHD D.K. supported by the National Institute Of Arthritis And Musculoskeletal And Skin Diseases of the National Institutes of Health under Award Number R01AR066124.

## Web resources

The URLs for data presented herein are as follows:

Exome Variant Server (NHLBI Exome Sequencing Project ESP6500): http://evs.gs.washington.edu/EVS/

ExAC: http://exac.broadinstitute.org

Human Genome Variation: http://www.hgvs.org/mutnomen/

Online Mendelian Inheritance in Man (OMIM): http://www.omim.org/ SeattleSeq: http://snp.gs.washington.edu/

**Figure 4.** Expression of *MYH3* in bone and cartilage. A) RNA was isolated from a 17-week old fetus and cDNA was generated using standard protocols^27^. *MYH3* primers were predicted to yield amplicons of 313 bp from genomic DNA (“Control”) and 206 bp from cDNA. *MYH3* cDNA expression was detected in bone thymus, placenta, heart, brain, and liver. Expression was not detected in cartilage, lung, and kidney. B) Quantitative PCR was used to assess the relative expression levels of *MYH3* cDNA in bone, cartilage, brain, heart, liver and lung. *MYH3* cDNA levels were detected in bone at a level similar to that observed in brain and heart and was barely detectable in lung and cartilage.

